# Negative memory engrams in the hippocampus enhance the susceptibility to chronic social defeat stress

**DOI:** 10.1101/379669

**Authors:** Tian Rui Zhang, Amanda Larosa, Marie-Eve Di Raddo, Vanessa Wong, Alice S. Wong, Tak Pan Wong

## Abstract

The hippocampus has been highly implicated in depression symptoms. Recent findings suggest that the expression and susceptibility of depression symptoms are related to the enhanced functioning of the hippocampus. We reasoned that hippocampal engrams, which represent ensembles of neurons with increased activity after memory formation, could underlie some contributions of the hippocampus to depression symptoms. Using the chronic social defeat stress (CSDS) model, we examined social defeat-related hippocampal engrams in mice that are either susceptible or resilient to the stressor. TetTag mice were used to label social defeat-related hippocampal ensembles by LacZ. Engram cells correspond to ensembles that were reactivated by the same stressor.

Compared to resilient and non-stressed control mice, we found that in both the dorsal and ventral hippocampal CA1 regions, susceptible mice exhibited a higher reactivation of social defeat-related LacZ-labeled cells (i.e. engram cells). The density of CA1 engram cells correlated with the level of social avoidance. Using DREADD to reactivate social defeat-related but not neutral contextual stimuli-related CA1 engram cells decreased social interaction. Increased engram cells in susceptible mice were region specific and could not be found in the dentate gyrus. Susceptible mice exhibited more negative stimuli-, but not neutral stimuli-, related CA1 engram cells than resilient mice in the dorsal hippocampus. Finally, chronic, but not a short and subthreshold, social defeat protocol was necessary to increase CA1 engram cell density. Together, our findings reveal that the susceptibility to CSDS is regulated by hippocampal CA1 engrams for negative memory. Hippocampal engrams for negative memory may underlie the vulnerability and expression of cognitive symptoms in depression.

**Significance statement:** We provided evidence that negative memory hippocampal engrams contribute to the susceptibility to developing depression-related behavior after chronic social defeat stress. The activation of positive memory engrams have been shown to alleviate depression-related behaviors, while our findings reveal the pathological roles of negative memory engrams that could lead to those behaviors. Increased negative memory engrams could be a downstream effect of the reported high hippocampal activity in animal models and patients with depression. Unlike affective symptoms, we know much less about the cellular mechanisms of the cognitive symptoms of depression. Given the crucial roles of hippocampal engrams in memory formation, enhanced reactivation of negative memory engrams could be an important cellular mechanism that underlies the cognitive symptoms of depression.

## Introduction

Apart from affective symptoms like sad mood, anhedonia, hopelessness and low self-esteem, cognitive symptoms are common in depression. A prominent cognitive symptom of depression is the negative bias in cognitive processing and memory formation (for review, see (Disner et al., 2011; Joormann and Quinn, 2014)). Depressed patients show enhanced encoding and recall of mood-congruent negative memory (Koster et al., 2010), less forgetting of negative memory (Hertel and Gerstle, 2003), and impaired recall of positive memory (Gaddy and Ingram, 2014). Rumination, which is related to repetitive recall of negative memory (Lyubomirsky et al., 1998), is also common in depression (Nolen-Hoeksema, 2000). Increasing findings suggest that bias in cognitive processing in depression could be associated with changes in the hippocampus, a brain region that is known for its role in memory formation (Squire, 1992).

The hippocampus has long been implicated in the manifestation of depression symptoms. Meta analyses have revealed reduced hippocampal volume in depressed patients (Videbech and Ravnkilde, 2004; McKinnon et al., 2009). Therapeutic effects of classical (e.g. fluoxetine) and fast-acting (e.g. ketamine) antidepressants have been associated with hippocampal neurogenesis (Santarelli et al., 2003) and altered hippocampal glutamate receptor function (Maeng et al., 2008; El Iskandrani et al., 2015), respectively. Recent findings suggest that increased hippocampal function could contribute to the biased cognitive processing in depression. Imaging studies revealed increased hippocampal responses to sad faces (Fu et al., 2004) and stronger hippocampus-amygdala connectivity during negative information encoding (Hamilton and Gotlib, 2008) in depressed patients. Moreover, attenuated hippocampal responses to negative stimuli could be induced by antidepressants (Mayberg et al., 2000; Fu et al., 2004) and observed in remitted depressed patients (Thomas et al., 2011). Using chronic social defeat stress (CSDS) as an animal model for depression-related behaviors (Krishnan et al., 2007), the expression of these behaviors has been associated with increased activity of the ventral hippocampal dentate gyrus (DG) region (Anacker et al., 2018). Suppressing ventral hippocampal glutamatergic inputs to the nucleus accumbens can enhance stress resilience of this model (Bagot et al., 2015). Increased hippocampal activity could affect the formation of engrams, which are ensembles of neurons that showed increased activity during memory formation and recall (Josselyn et al., 2015; Tonegawa et al., 2015). Hippocampal engrams have been associated with the expression of depression-related behaviors, so that reactivating positive memory-related hippocampal engrams can induce antidepressant effects (Ramirez et al., 2015). Increased hippocampal activity could also facilitate the formation and enhance the activity of negative memory engrams. Whether negative memory engrams contribute to the expression of depression-related behaviors warrants further investigations.

In the current study, we investigated the formation and activity of social defeat-related hippocampal engrams in mice that were stressed under the CSDS paradigm/protocol. We used TetTag mice to tag hippocampal neurons that were activated by social defeat with LacZ (Reijmers et al., 2007). The reactivation of LacZ labeled cells caused by the same stressor, which represent engram cells, was also examined. The CSDS model allows us to separate mice according to their individual differences in stress susceptibility. We found that mice that were susceptible to CSDS had more social defeat-related engram cells in the hippocampal CA1 region than non-stressed control mice and mice that were resilient to this stressor.

## Materials and Methods

### Animals

Male TetTag mice were obtained from the *Jackson Laboratory* (stock no. 008344, (Reijmers et al., 2007)). Bi-transgenic TetTag mice with a C57 background carry a cFos-driven tetracycline-controlled transactivator (tTA) protein construct and a tetracycline-responsive regulatory element (tetO) driven beta-galactosidase (LacZ) construct. The cFos promoter can be activated by neuronal activity. This strain has been used for labeling activated neurons by the expression of LacZ via a doxycycline off (Dox-off) mechanism as previously described (Reijmers et al., 2007). Double hemizygote TetTag mice were bred with wild type C57 mice (*Charles River*). Only male double hemizygote offspring (approximately 1/8 of all offspring) were used in this study. Breeding pairs and offspring were fed with Dox-containing food (40 mg/kg, *Envigo*) *ad libitum* in a 12 hour light/dark cycle (light on from 8AM to 8PM). LacZ labeling can be induced by feeding TetTag mice with Dox-free food (Dox off), which allows the cFos-driven expression of tetracycline transactivator (tTA) to activate the tetO-LacZ construct. The activation of tetO during Dox off also triggered the expression of a tetracycline-insensitive tTA (with a H100Y point mutation), which sustained the expression of LacZ even after the reintroduction of Dox to maintain long-term labeling of activated neurons. The average age of the mice was 3 months. Offspring of TetTag mice that expressed only the cFos-tTA construct were used in the DREADD experiment (see below). Finally, male retired breeders of the CD1 strain (*Charles River*) were used for defeating mice of C57 strains. All experiments were approved by the Facility Animal Care Committee at Douglas Institute and followed the guidelines from Canadian Council on Animal Care (protocol no.: 2010-5935).

### Chronic Social Defeat Stress (CSDS)

TetTag mice were defeated by male retired breeders of the CD1 strain during social defeat. Resident CD1 mice were housed in a partitioned compartment of a rat cage, like the type of cage used for habituation, before social defeat. Each CD1 mouse was screened for its aggressiveness by attacking intruders and only those with a < 60 seconds latency were selected. The CSDS paradigm consisted of 8 episodes of defeat. In each defeat episode, a CD1 mouse was allowed to attack a TetTag mouse for up to 12 attacks in a maximum period of 5 minutes. Following each social defeat episode, each TetTag mouse was housed next to the CD1 mouse in the neighboring compartment separated by a perforated partition for 24 hours. Without physical contacts, TetTag mice were stressed during cohousing by the presence of visual and odor stimuli from the CD1 mouse. Each TetTag mouse was paired with a new CD1 mouse in each of the 8 episodes of social defeat to prevent reduced number of attacks due to repeated cohousing. Control non-stressed mice, which were only handled and weighted daily, were pair-housed in neighbor partitions in a rat cage for 8 days. After 8 daily episodes of defeat or pair-housing, social behavior of stressed and control mice were examined by a social interaction (SI) test.

### Social Interaction (SI) Test

The SI test consisted of two 150 seconds long sessions of exploration in a Plexiglas open field (44 cm x 44 cm). An empty perforated enclosure (10 cm x 5 cm x 30 cm) was placed in the center of the north side of the open field during the first open field session. After the end of the first open field session, a CD1 mouse was put into the enclosure before the second open field session began. Both open field sessions were performed under ambient red light, with static white noise at 60 dB. Time spent in the interaction zone (10 cm around the enclosure) during the first (empty) and second (with a CD1 mouse) open field sessions were estimated from recorded videos of these sessions using the software TopScan LITE (*Clever system Inc*.). The SI ratio was calculated by dividing the time mice spent in the interaction zone in the second open field session with the time they spent in the interaction zone in the first open field session. We also measured time TetTag mice spent in the two corners zones (10 cm x 10 cm) on the opposite side of the enclosure, which were farthest away from the enclosure. Corner ratios were calculated by dividing time TetTag mice spent in those corners in the second open field session by the time they spent there in the first session. Susceptible mice were defined as animals having a social interaction ratio of less than 1, indicating they spent less time in the interaction zone when a CD1 mouse was present. Resilient mice were defined having a social interaction ratio of greater than 1 and spent at least 50 seconds in the social interaction zone during the second open field session.

### Ensembles reactivation

After the SI test, both stressed and control mice were housed singly in mouse cages. To reactivate ensembles that were related to social defeat in stressed mice, we gave stressed mice an extra episode of social defeat, followed by co-housing in the neighbor compartment with a CD1 mouse in a partitioned rat cage for 90 minutes to trigger cFos expression. Ensembles related to contextual information of the rat cage were reactivated in control mice to express cFos by co-housing them with another control mouse in neighbor compartments of a partitioned rat cage for 90 minutes. After ensemble reactivation, mice were anesthetized and perfused by heparin-containing phosphate-buffered saline (PBS) and 4% paraformaldehyde solution (PFA). Brains were extracted from the skulls, postfixed in PFA overnight and cryoprotected in 30% sucrose-containing PBS.

### Immunohistochemistry

Fixed brains were snap-frozen in dry ice-chilled isopentane before being cut into 35 μm-thick sections using a cryostat (*Leica*). Brain sections were washed with PBS (five 5-minute washes; a similar washing procedure was used between all antibody incubations), followed by a 30 minutes incubation in 0.3% NaBH_4_ (*Sigma*) to quench endogenous fluorescence. After PBS washes, sections were incubated for one hour in a blocking solution (3% normal goat serum and 0.1% Triton in PBS (PBS-T), this blocking solution was also used for diluting antibodies). For triple immunofluorescent staining, sections were incubated overnight at 4°C with the first primary antibody (mouse monoclonal LacZ antibody, 1:2000 (*MP Biomedicals*, 08633651)). The next day sections were washed by PBS-T and incubated with the first secondary antibody (donkey anti-mouse Alexa 674 antibody, 1:2000 (*Abcam*, Ab150107)) for three hours at room temperature. In the same fashion, incubations were done for the second primary antibody (rabbit polyclonal cFos antibody, 1:40,000 (*Sigma*, F137)) and the corresponding secondary antibody (goat antirabbit Alexa 488 antibody, 1:4000 (*Life Technologies*, A11034)). Finally, sections were incubated with 600 nM 4’,6-Diamidino-2-Phenylindole (DAPI) (*Life Technologies*, D3571) for 10 minutes. Triple-labeled sections were mounted on a slide, covered with VectaShield anti-fade mounting medium (*Vector Laboratories*) and sealed with nail polish. The stained sections were scanned using a slide scanner (*Olympus* VS120) with the VS-ASW acquisition software to a magnification of 20X with eleven 15 μm-thick z-sections. Sections were stitched together by the VS-ASW software (*Olympus*).

For 3,3’diaminobenzidine (DAB, *Sigma*) staining of cFos, after primary antibody incubation and washes, slices were incubated for 1 hour with a biotinylated goat anti-rabbit antibody (1:500, *Vector Laboratories*, BA-1000), followed by an hour long incubation with the ABC reagent (1:250, *Vector Laboratories*, PK-7200). Sections were finally incubated with DAB (0.6%) and H2O2 for 2 minutes to visualize staining.

### Cell counting

Analysis of the digital slides from the slide scanner were done manually with the help of Fiji (*ImageJ*). As there are regional differences in inputs, projections and functions between dorsal and ventral and hippocampus (Fanselow and Dong, 2010), cell counting was performed in both regions. For CA1 counting, a 400 μm (width) by 200 μm (height) counting window was used for counting LacZ-, cFos- and DAPI-labeled cells in the dorsal and ventral hippocampus. Density of single (LacZ or cFos)- and double (LacZ and cFos)-labeled cells were estimated by dividing their numbers with the number of DAPI-labeled cells in the counting window. Only neurons in the *stratum pyramidale* were counted. Since we found no LacZ cells in the pyramidal layer of the CA3 region in both control and stressed mice, this hippocampal region was excluded from further analysis. Finally, for cell counting in the dentate gyrus (DG) region, due to the low number of double labeled cells in the DG, the entire DG granule cell layer in the dorsal and ventral hippocampus in each section was counted. To compare the density of DAPI cells between animal groups, we controlled for the differences in the size of DG between sections by normalizing the density of DAPI cells by the length of the granule cell layer. Three to five sections from each hippocampal region of each mouse were used for counting. Data from these sections were averaged and only the mean densities of single and double labeled cells of each mouse were used for statistical analysis.

### Experimental design and statistical analysis

All statistical analyses were performed using GraphPad Prism 7. Normality of data was examined by the Shapiro-Wilk’s test. All data were presented as mean ± SEM.

#### Experiment 1: Social defeat-related hippocampal engrams in mice with different stress susceptibility

Adult male TetTag mice were off Dox for 4 days during habituation before being stressed by 8 episodes of social defeat (Figure 1A). Habituation has been shown to reduce the labeling of hippocampal neurons from being housed in a novel environment (Radulovic et al., 1998). During habituation, two TetTag mice were housed in neighboring compartments of a rat cage. We found that 2 social defeat episodes were sufficient to induce cFos (Figure 1B) and LacZ expression (Figure 1C) in the hippocampus. LacZ labeling was therefore stopped after 2 episodes of social defeat by Dox (1 g/kg) for 1 day, followed by regular Dox food (40 mg/kg) to prevent further LacZ labeling. Note that there was no neuronal LacZ expression in the hippocampus of TetTag mice that were always on Dox (Figure 1C). After a total 8 episodes of social defeat, TetTag mice were examined by the SI test. One day after the SI test, social defeat-related ensembles of stressed mice were reactivated by another defeat episode. Control TetTag mice were treated similarly as stressed mice, but they were only handled after habituation and during ensemble reactivation. Mice were sacrificed 90 minutes after ensemble reactivation for immunostaining. There were 29 stressed and 9 control mice in this experiment. Stressed mice were furthered divided into susceptible and resilient mice according to their SI ratio (Figure 2). Due to the distinct roles of the dorsal and ventral hippocampus in spatial and emotional functions, data from dorsal and ventral hippocampus were separately compared. Density of LacZ, cFos, engram and DAPI cells were separately compared between the three mouse groups using one-way ANOVA and post hoc Tukey’s test (Figure 4 and 7).

**Figure 1:**
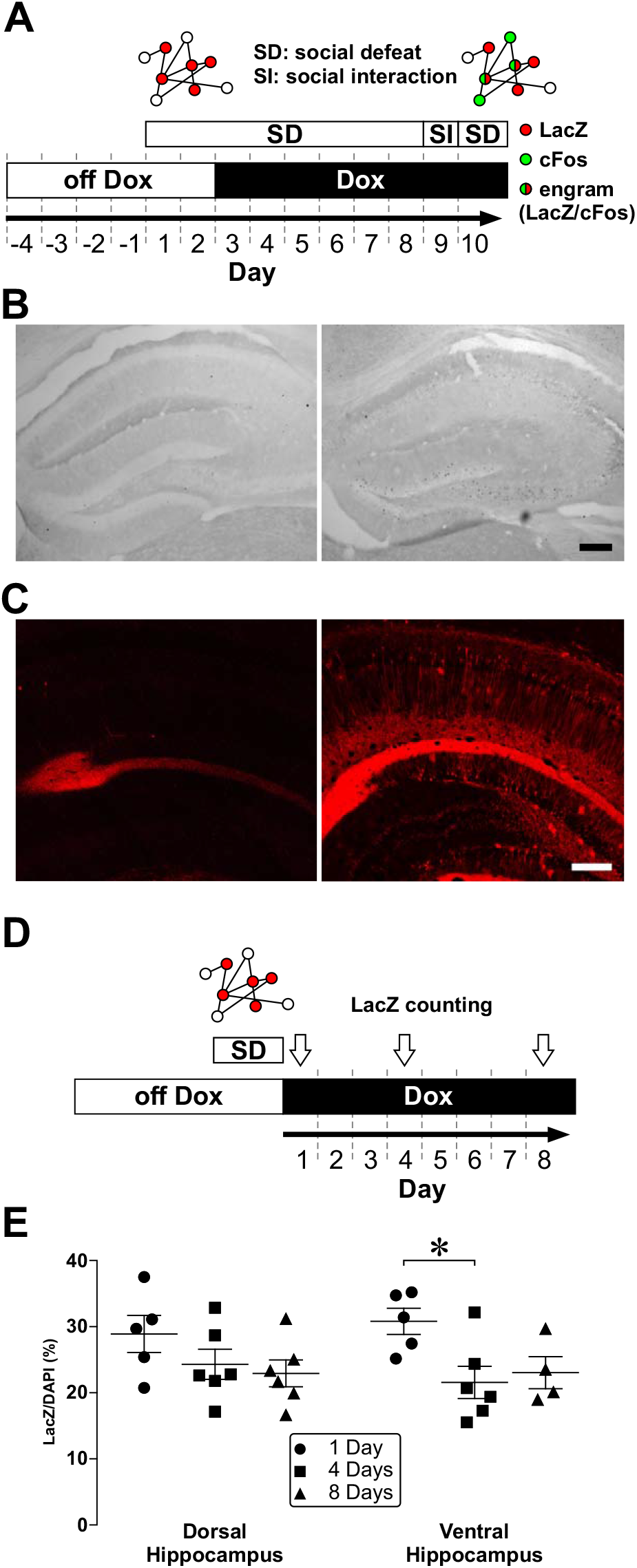
*Social defeat triggers the formation of hippocampal ensembles*. (**A**) A schematic diagram of the experimental design. TetTag mice were off Dox for 4 days. After two episodes of social defeat (SD) on day 1 and 2, labeling was blocked by putting mice on Dox-containing food. Mice were then stressed by 6 more episodes of SD. The interaction between TetTag mice and aggressors of the CD1 strain was examined in a social (SI) interaction test. One day after the SI test, mice underwent one more episode of SD to trigger ensembles reactivation. Mice were sacrificed 90 minutes after the last episode of SD. Cartoons above the experimental plan depict the labeling of activated neurons during the first two episodes of CSDS (red, LacZ), during the last episode of SD (green, cFos), and engram cells that expressed both signals (red/green). (**B**) cFos stained dorsal hippocampal sections from a control mouse that was housed in its home cage (*Left*). (*Right*) cFos stained dorsal hippocampal sections from another control mouse that was defeated by a single episode of SD one day earlier. Scale bar = 200 μm. (**C**) LacZ staining of ventral hippocampal neurons from TetTag mice that were off doxycycline-containing food during labeling (Dox off, *right*). A stained section from a mouse that was on doxycycline-containing food during labeling was shown on the *Left* (Dox on). Note that apart from nonspecific staining near the hippocampal fissure, LacZ cells and processes cannot be found in tissue from the Dox on mouse. Scale bar = 200 μm. (**D**) A schematic diagram of the experimental design for testing the stability of LacZ expression after Dox on. TetTag mice were off Dox for 4 days. After two episodes of social defeat (SD), labeling was blocked by putting mice on Dox. TetTag mice were sacrificed 1 day, 4 days and 10 days later (white arrows). (**E**) Scatter plots summarize the density of LacZ positive neurons in the CA1 region of the dorsal and ventral hippocampus of TetTag mice at different time points after labeling. * p < 0.05, post hoc Tukey’s test vs. data from day 3 group in each hippocampal region after ANOVA.

**Figure 2:**
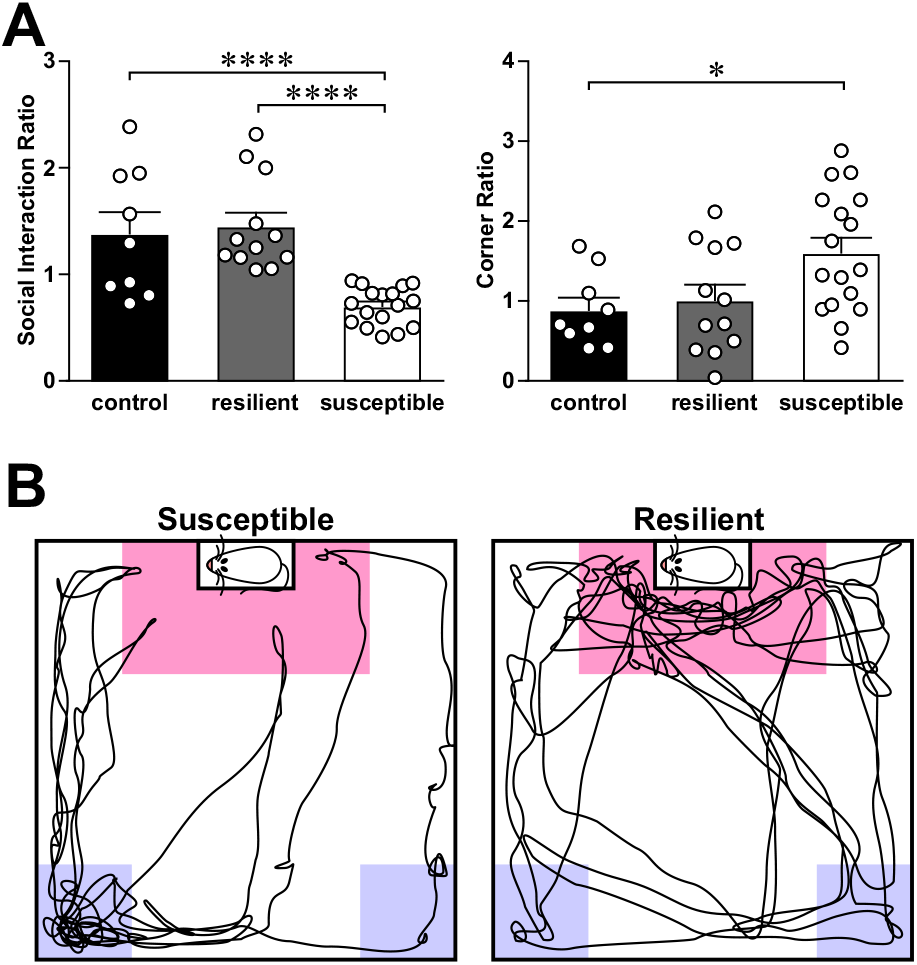
*Susceptible but not resilient mice expressed social avoidance after chronic social defeat stress*. (**A**) Histograms summarize the social interaction ratio (*Left*) and the corner ratio (*Right*) of susceptible, resilient and non-stressed control mice. * p < 0.05, **** p < 0.0001, post hoc Tukey’s test after ANOVA. (**B**) Example tracks of a susceptible (*Left*) and a resilient mouse during the second open field session of the SI test. The pink and purple zones are the virtual interaction and corner zones, respectively. Note the cluster of tracks in the interaction and corner zones for the resilient and susceptible mice, respectively.

#### Experiment 2: Impact of activating social defeat-related engrams on social interaction

Adult male cFos-tTA mice were on Dox while they were bilaterally injected into the dorsal hippocampus with 0.5 μl AAV-PTRE-tight-hM3Dq-mCherry (Zhang et al., 2015). One week after virus injection, they were off Dox while they were habituated in pairs in partitioned rat cages. Some mice (n = 18) were stressed by a short and subthreshold social defeat protocol, which consists of only 2 episodes of social defeat described in *Experiment 1*. Other mice (n = 13) served as controls and were only weighed and handled daily for 2 days. One day after defeat or handling, mice were examined by the SI test. One hour before the SI test, stressed and control mice were randomly selected to receive i.p. injection of either saline or clozapine-N oxide (3 mg/kg). Mice were sacrificed and perfused after the SI test to examine virus expression. SI ratios from these four mouse groups were compared using one-way ANOVA and post hoc Tukey’s test (Figure 6).

#### Experiment 3: Neutral contextual stimuli-related hippocampal engrams in mice with different stress susceptibility

To label neutral contextual stimuli-related engrams, adult male TetTag mice were off Dox for 6 days during habituation before being stressed by 8 episodes of social defeat (Figure 8A). LacZ expression was arrested one day before social defeat by 1 g/kg Dox, followed by 40 mg/kg Dox throughout the 8 episodes of social defeat. One day after the last episode of defeat, TetTag mice were examined by the SI test, followed one day later with the reactivation of ensembles by another episode of social defeat. Control TetTag mice were treated similarly as stressed mice, but were only handled after habituation and during ensemble reactivation. Mice were sacrificed 90 minutes after ensemble reactivation for immunostaining. We had a total 34 stressed and 19 control mice. Stressed mice were furthered divided into susceptible and resilient mice according to their SI ratio (Figure 2). Due to the distinct roles of the dorsal and ventral hippocampus in spatial and emotional functions, data from the dorsal and ventral hippocampus were separately compared. Density of LacZ, cFos, engram and DAPI cells were separately compared between the three mouse groups using one-way ANOVA and post hoc Tukey’s test (Figure 8).

#### Experiment 4: Impact of subthreshold social defeat on hippocampal engrams

Adult male TetTag mice were off Dox for 4 days during habituation before being stressed by 2 episodes of social defeat (Figure 9A). LacZ labeling was stopped by putting mice on Dox after the second social defeat episode. Stressed mice were examined by the SI test either 1 day (n = 8; defeated no delay) or 7 days (n = 9; defeated with delay) after social defeat, followed by the reactivation of ensembles one day after the SI test by another social defeat episode. Control TetTag mice (n = 8) were treated similarly as stressed mice but were only weighed and handled after habituation and during ensemble reactivation. Mice were sacrificed 90 minutes after ensembles reactivation for immunostaining. Data from the dorsal and ventral hippocampus were separately compared. Density of LacZ, cFos, engram and DAPI cells were separately compared between the three mouse groups using one-way ANOVA and post hoc Tukey’s test (Figure 9).

## Results

Using the CSDS protocol (Figure 1A), we identified 17 susceptible mice that displayed social avoidance (i.e. social interaction ratio (SI ratio) < 1) and 12 resilient mice that showed normal social behavior after stress (Figure 2A, B). In addition, 9 control nonstressed mice were habituated and fed with Dox and normal food like the stressed mice. These mice were pair-housed with another TetTag or nontransgenic littermates for 8 days after habituation and were only handled daily. One-way ANOVA revealed a significant difference in SI ratio between animal groups (F(2,35) = 15.7; p = 1.37E-05) with susceptible mice having lower SI than control (post-hoc Tukey’s test: p = 5.24E-04) and resilient mice (p = 4.28E-05).

We also examined the time mice spent in the corners of the open field during the SI tests (Figure 2A, B). Susceptible mice spent significantly more time in corner zones than control and resilient mice in the second open field session when a social object was present in the enclosure. Comparing the corner ratios revealed a significant between group difference (F(2,35) = 4.56; p = 0.0173). The corner ratio of susceptible mice was significantly higher than control (p = 0.0319). Although susceptible and resilient mice displayed distinct behaviors during the SI test, we did not observed differences in the number of attacks (11.8 ± 0.2 for susceptible mice vs. 11.5 ± 0.3 for resilient mice) and the duration of social defeat (i.e. time used for all attacks: 153.4 ± 13.0 seconds for susceptible mice vs. 143.6 ± 13.9 seconds for resilient mice) between these two groups. Finally, all three mouse groups showed similar weight gain from Day 1 to Day 8 after habituation (1.44 ± 0.30 g for control mice; 1.85 ± 0.34 g for resilient mice; 1.38 ± 0.39 g for susceptible mice).

### Susceptible mice displayed more engram cells in the hippocampal CA1 region than resilient and control mice

To find out if stress susceptibility is related to the reactivation of hippocampal ensembles that were labeled during social defeat, we examined the reactivation of LacZ ensembles that were formed during the first 2 episodes of social defeat by an extra episode of social defeat one day after the SI test in stressed mice (see Figure 1A). After 90 minutes following the extra episode of social defeat, stressed mice were sacrificed for immunostaining of LacZ and cFos to reveal activated ensembles. Control mice were only exposed to the context during LacZ expression (i.e. pair-housed in a partitioned rat cage) for 90 minutes to examine the reactivation of neutral context-related LacZ ensembles. Reactivated engram cells in ensembles were represented by double labeled cells that expressed both LacZ and cFos (Figure 3). The formation of these double labeled engram cells cannot be explained by probabilistic reasons, since the density of double labeled cells in all mouse groups was significantly higher than chance (LacZ/DAPI x cFos/DAPI) in both the dorsal (control: t(8) = 4.23, p = 0.003; resilient: t(11) = 7.63, p = 1.03E-05, susceptible: t(15) = 7.45, p = 2.06E-05) and the ventral hippocampus (control: t(8) = 5.10, p = 9.35E-04; resilient: t(11) = 3.87, p = 0.003, susceptible: t(15) = 6.62, p = 8.13E-06).

Densities of LacZ, cFos and engram cells in the dorsal and ventral hippocampus were separately compared in order to reveal region specific differences. Although only resilient and susceptible mice were stressed by CSDS, we did not observe significant differences in the density of LacZ (Figure 4A) and cFos cells (Figure 4B) between the three mouse groups in both the dorsal and ventral hippocampal CA1 regions. However, we found that the density of engram cells in susceptible mice was significantly higher than control and resilient mice in both the dorsal (Figure 4C, F(2,35) = 18.4; p = 3.54E-06; post-hoc Tukey’s test: control vs. susceptible, p = 8.88E-05, resilient vs. susceptible, p = 2.21E-05) and the ventral hippocampus (F(2,35) = 16.2; p = 1.07E-05; post-hoc Tukey’s test: control vs. susceptible, p = 1.80E-03, resilient vs. susceptible, p = 1.52E-05). Since we have previously shown that CSDS has different impacts on hippocampal volume in susceptible and resilient mice (Tse et al., 2014), we asked if changes in the density of CA1 neurons were responsible for the increase in engram cell density in susceptible mice (Figure 4D). However, we did not observe differences in the density of DAPI CA1 cells between these mouse groups.

Although we did not observe significant changes in the density of LacZ cells between the three animal groups, when we analyzed data from the dorsal and ventral hippocampus separately, two-way ANOVA analysis of the effect of dorsal and ventral regions and the animal group on the density of LacZ cells revealed a significant effect of animal group (Effect of animal groups: F(2,70) = 4.23, p = 0.0185), and a significantly higher LacZ cell density in susceptible mice than in resilient mice when both dorsal and ventral data were pooled together (post-hoc Tukey’s test: control vs. susceptible, p = 0.121, resilient vs. susceptible, p = 0.025). This slight but significant increase in LacZ cell density may underlie the increased engram cell formation in susceptible mice. To test this, we normalized the engram cell density with the density of LacZ cells and compared the data between the 3 animal groups. After normalization, susceptible mice still have more engram cells than both control and resilient mice in the dorsal hippocampus (F(2,35) = 9.06; p = 6.72E-04; post-hoc Tukey’s test: control vs. susceptible, p = 0.00256, resilient vs. susceptible, p = 4.23E-03). Susceptible mice also have more normalized engram cells than resilient mice in the ventral hippocampus (F(2,35) = 6.78; p = 6.72E-04; post-hoc Tukey’s test: control vs. susceptible, p = 0.0738, resilient vs. susceptible, p = 3.21E-03). These findings strongly suggest that susceptible mice have more social defeat-related CA1 engram cells in both the dorsal and ventral hippocampus than resilient and control mice.

The higher engram cell density in susceptible mice than resilient and control mice suggest that engram cell density is related to the expression of depression-related behavior of these mice. Indeed, we found that CA1 engram cell density in both the dorsal (Figure 5A, R^2^ = 0.192, p = 5.98E-03) and ventral hippocampus (R^2^ = 0.166, p = 0.0110) of all tested mice correlated negatively with the SI ratio. When we examined the relationship between CA1 engram cell density and the corner ratio, we also found a significant correlation between dorsal CA1 engram cell density and corner ratios (Figure 5B, R^2^ = 0.176, p = 8.65E-03). However, the correlation between ventral CA1 engram cell density and corner ratios did not reach a significant level (R^2^ = 0.0627, p = 0.129). These findings suggest that high CA1 engram cell density in susceptible mice is related to the expression of social avoidance.

Findings from the correlation analyses suggest that reactivating CA1 engram cells can trigger social avoidance. To test that, we used the cFos-tTA offspring from TetTag mice that lack the LacZ construct (Figure 6A). While these mice were fed with Dox, we bilaterally injected AAV-PTRE-tight-hM3Dq-mCherry into the dorsal hippocampi of adult cFos-tTA mice (Figure 6B). After one week, we put these mice off Dox for two days before stressing mice with a subthreshold social defeat protocol with only 2 episodes of defeat. tTA from activated neurons will bind to TRE to trigger the expression of excitatory DREADD hM3Dq in these neurons, which can be activated by a DREADD ligand clozapine N-oxide (CNO). cFos-tTA mice that have received AAV-PTRE-tight-hM3Dq-mCherry injection but no social defeat served as controls. One day after the last social defeat episode, we injected stressed and control mice with either vehicle or CNO (3 mg/kg) at 1 hour before the SI test. We observed significant changes in SI ratio between the 4 groups of mice (F(3,27) = 4.22; p = 6.72E-04). Pairwise comparisons revealed a significant difference between the control CNO and the defeated CNO groups (post-hoc Tukey’s test: control CNO vs. defeated CNO, p = 7.36E-03), suggesting the activation of social defeat-related hippocampal CA1 engrams reduces social interaction.

### CSDS reduced engram cell density in the hippocampal dentate gyrus region

Fear memory formation and recall have been associated with engrams in the DG (Liu et al., 2012; Deng et al., 2013; Denny et al., 2014). We next examined if susceptible mice also express more DG engram cells than other mouse groups. Similar to findings we observed from the CA1 region, we did not find changes in the density of LacZ (Figure 7A) and cFos cells (Figure 7B) in the DG between the three mouse groups. Interestingly, we saw a trend that engram cell density in control mice to be higher than both resilient and susceptible mice in the ventral DG (Figure 7C, F(2,34) = 3.21; p = 0.0529). The fact that both the resilient and susceptible groups displayed similar changes in engram cell density suggests an effect due to stress. Similar to our prediction, engram cell density in the ventral DG of control mice remained higher than pooled data from the susceptible and resilient groups (control vs. stressed mice: t(35) = 3.33, p = 2.04E-03). Similarly, in the dorsal hippocampus, we observed a trend where engram cell density in stressed mice is lower than control mice (control vs. stressed mice: t(36) = 1.86, p = 0.0717). Finally, we compared the density of DAPI-labeled neurons in the DG of the three mouse groups and observed no difference between groups (Figure 7D), even after data of susceptible and resilient mice were pooled together. DG engram cells therefore may not contribute to the susceptibility to CSDS.

### Engram cell formation in susceptible mice caused by neutral stimuli

Even after habituation, we saw overlapping LacZ and cFos ensembles in control non-stressed mice. The density of engram cells was higher than the chance levels (i.e. LacZ/DAPI vs. LacZ/DAPI x cFos/DAPI; dorsal hippocampus: t(8) = 4.23, p = 0.003; ventral hippocampus: t(8) = 5.10, p = 9.35E-04). These findings suggested that during Dox off, the exposure to neutral contextual information triggered the formation of LacZ ensembles in the CA1 region. These ensembles were reactivated by re-exposure to the same context. Engram cells observed in stressed mice in Figure 4 were likely due to the reactivation of ensembles that are related to both neutral (contextual information) and negative (social defeat) stimuli. To find out whether susceptible mice also exhibited higher reactivation of neutral stimuli-related LacZ ensembles than other mouse groups, we stopped LacZ labeling before social defeat and studied ensembles reactivation (Figure 8A). Both control and stressed mice were habituated off Dox in the partitioned rat cage for 6 days to maintain a similar duration of LacZ expression as in previous experiments (see Figure 4, 4 days habitation plus 2 days of social defeat). LacZ expression was stopped one day before social defeat by Dox, followed by similar procedures we used for CSDS and SI test in Figure 1A (Figure 8A). We identified 13 susceptible mice and 5 resilient mice in this experiment. Compared with control mice (n = 10), we again did not observe differences in the density of LacZ (Figure 8B) and cFos cells (Figure 8C) between the 3 animal groups. Interestingly, we also did not find significant differences in the density of engram cells in the dorsal hippocampus between the animal groups. However, we found that susceptible mice expressed more engram cells in the ventral CA1 region than control mice (Figure 8D, F(2,22) = 5.05; p = 0.0156; post-hoc Tukey’s test: control vs. susceptible, p = 0.0259). When we compared the density of DAPI neurons in the CA1 of the three mouse groups, we found no difference between-groups (Figure 8E), suggesting no changes in neuronal density. Thus, enhanced engram cell formation in the dorsal hippocampus of susceptible mice was largely limited to those that respond to negative-but not neutral stimuli.

Exposure to chronic, but not acute, stressors is crucial for the development of depression-related behaviors (McGonagle and Kessler, 1990; McEwen, 2004). Indeed, we showed that stressing mice with only 2 episodes of social defeat did not result in social avoidance (Figure 6). To find out if this subthreshold number of defeat episodes is too weak to induce engram cell formation, we examined LacZ, cFos, and engram cell density in non-stressed mice and mice that were stressed by 2 episodes of social defeat (Figure 9A). In addition to examining the SI ratio of stressed mice 1 day after social defeat (no delay), we examined SI ratio of another group of mice that were stressed by the subthreshold protocol 7 days after social defeat (with delay), which corresponds to the time point when engram cells were examined in other experiments (see Figure 1A). This control group could reveal whether engram cell density remains stable under delayed observation. Similar to what we showed in Figure 6, this subthreshold social defeat paradigm did not result in social avoidance if SI was examined 1 day after defeat (no delay). Surprisingly, when SI was examined 7 days after defeat (with delay), we found that defeated mice exhibited social avoidance (Figure 9A, F(2,25) = 5.68, p = 9.29E-03; post-hoc Tukey’s test: control vs. defeated with delay, p = 0.0194, defeated no delay vs. defeated with delay, p = 0.0221). Since responses to aversive experience tend to generalize with time, we reasoned that the expression of social avoidance in defeated with delay mice was due to the spread of fear to nonspecific avoidance during the SI test. When we compared ensemble activity in these mice, we found that density of LacZ cells in both the dorsal and ventral hippocampus was significantly higher in defeated no delay mice than other mouse groups (Figure 9B. Dorsal hippocampus: F(2,21) = 11.4, p = 4.50E-04; post-hoc Tukey’s test: control vs. defeated no delay, p = 6.77E-04, defeated no delay vs. defeated with delay, p = 3.27E-03. Ventral hippocampus: F(2,22) = 6.18, p = 7.41E-03; post-hoc Tukey’s test: control vs. defeated no delay, p = 7.09E-03, defeated no delay vs. defeated with delay, p = 0.0475). The increase in LacZ cell density one day after defeat revealed the activation of CA1 neurons induced by this stressor. The decrease in LacZ cell density caused by the delay may be due to the dissipation of LacZ signals we showed in Figure 1D. Unlike these changes in LacZ cells, we did not observe changes in the density of cFos (Figure 9C), engram (Figure 9D), and DAPI cells (Figure 9E) between these three groups. Our findings suggest that a subthreshold social defeat paradigm does not increase the density of CA1 engram cells in the hippocampus. The expression of social avoidance after a delay from subthreshold social defeat, possibly due to generalization, is likely related to non-CA1 mechanisms.

## Discussion

Our findings suggest that social defeat-related negative memory engrams in the hippocampal CA1 region are closely related to the expression of social avoidance in mice that are susceptible to CSDS. We found that susceptible, but not resilient, mice exhibited a higher density of CA1 engram cells than non-stressed control mice. Social avoidance not only correlated with the density of CA1 engram cells, but also was facilitated by activating social defeat-related dorsal CA1 engram cells using a DREADD approach. Finally, a subthreshold social defeat protocol that failed to induce social avoidance did not increase engram cell density. Taken together, our findings suggest that the reactivation of stress-related negative memory engram cells in the CA1 region contributes to the susceptibility to CSDS.

Using social defeat to induce the labeling (LacZ by the first 2 defeat episodes) and the reactivation (cFos by an extra defeat episode one day after the SI test) of hippocampal engrams, susceptible mice showed higher CA1 engram cell reactivation than resilient and control mice. Previous findings strongly suggest that the reactivation of CA1 engram cells are related to memory retrieval that was triggered by contextual information (Deng et al., 2013; Cai et al., 2016; Roy et al., 2017). Higher engram cell density in susceptible mice therefore may be due to the facilitated retrieval of memory that is related to the defeat experience. Contextual information related to social defeat in the SI test, such as the presence of a CD1 aggressor, may be sufficient to reactivate social defeat-related engrams in susceptible mice to enhance avoidance behavior. Alternatively, resilient mice may be able to cope with CSDS by suppressing negative memory engrams reactivation. Retrieval of mood congruent memory, which is commonly found in depression, has been suggested to reduce the ability of depressed patients for problem solving and sparing attention to positive information and memory (Conway and Pleydell-Pearce, 2000). Persistent recall of negative memory underlies rumination, which has been associated with the vulnerability to depression and the severity of depression symptoms (Alloy et al., 1999; Rude et al., 2003; Abela and Hankin, 2011) and perhaps most importantly, hippocampal activation (Denson et al., 2009; Mandell et al., 2014). Resilient mice may be able to suppress negative memory engram reactivation through a top down inhibitory control from the frontal lobe (Disner et al., 2011; Kircanski et al., 2012).

Susceptible mice may exhibit a bias of forming CA1 engram for negative stimuli. Compared to resilient mice, susceptible mice show higher engram cell density when they were exposed to negative but not neutral stimuli (Figure 8). This bias in engram formation cannot be found in the ventral hippocampus, where enhanced neutral- and negative-stimuli related engrams were found in susceptible mice. Ventral hippocampal activation has been proposed to underlie the susceptibility to CSDS, so that enhanced and suppressed ventral hippocampal activity could confer to the expression of susceptibility phenotypes (Anacker et al., 2018) and stress resilience (Bagot et al., 2015), respectively. Our findings further suggest that context-related reactivation of hippocampal engram cells contributes to stress susceptibility. Recent studies showed that defeated mice exhibited lower levels of avoidance if an anesthetized aggressive mouse or a non-CD1 strain mouse was used in a social interaction test (Krishnan et al., 2007; Venzala et al., 2012), supporting the idea that cognitive factors, such as social defeat-related contextual information, as important for the expression of social avoidance. Roles of the dorsal hippocampus in stress susceptibility may be overlooked when overall activity, but not context-related reactivation, of this region was examined.

Our findings that subthreshold social defeat failed to increase hippocampal CA1 engram cell density could have several implications. We found that stressed mice at one day after the subthreshold defeat protocol exhibited a significant increase in CA1 LacZ cell density, but no change in the SI ratio (Figure 9, the defeated no delay group), suggesting that high CA1 neuronal activation in both the dorsal and ventral hippocampus alone is not sufficient to induce social avoidance. When social behavior of a separated group of mice stressed by subthreshold social defeat was examined a week after defeat, we observed social avoidance but no change in the density of CA1 LacZ, cFos and engram cells. Given the well-known effect of time to generalize fear memory (Wiltgen and Silva, 2007), the delayed expression of social avoidance in mice stressed by subthreshold social defeat may be due to fear generalization. The lack of changes in the CA1 region in these mice suggests that brain regions that are related to fear generalization such as the DG (Yokoyama and Matsuo, 2016), amygdala (Botta et al., 2015), nucleus reuniens and PFC (Xu and Sudhof, 2013) may be responsible. Finally, our findings suggest that a chronic stressor, which is closely associated to the etiology of depression (McGonagle and Kessler, 1990; McEwen, 2004), rather than a short subthreshold stressor, is needed to induce changes in engram cells. Persistent plastic changes at the synaptic and cellular levels may be induced by repeated stress exposure to facilitate the reactivation of engram cells in susceptible mice.

Unlike the CA1 region, we did not find differences in DG engrams between susceptible and resilient mice. Instead, we observed lower engram cell density in stressed mice when compared to non-stressed control mice. DG plays important roles in pattern separation and is likely sensitive to changes in context (Leutgeb et al., 2007). Using TetTag mice, it has been shown that while engram cells were formed in both the CA1 and DG during contextual learning, subsequent exposure to the same context favored the reactivation of CA1, but not DG engram cells (Deng et al., 2013). Since mice were kept in a similar context for multiple days during CSDS, CA1 instead of DG ensembles may be preferentially reactivated under this behavioral paradigm. Indeed, compared to 4-9% reactivation of DG cells in a relatively short behavioral task such as fear conditioning (Liu et al., 2012; Denny et al., 2014; Stefanelli et al., 2016), only ~0.5% of DG cells were reactivated by social defeat in the current study. A recent study however has shown that CSDS increased the number of cFos cells in the ventral, but not the dorsal DG (Anacker et al., 2018). The difference between findings from this study and the current study may be due to the use of a stronger defeat paradigm (5-minute-long vs. 3-minute-long defeat in the current study) and different ways of quantification (number of cFos cells per hemisphere vs. normalized cFos cell density by DAPI in the current study).

Similar to a recent report using the same mouse model (Deng et al., 2013), we were not able to detect CA3 engram cells in TetTag mice due to low LacZ expression in this hippocampal region (Figure 3). In the experiment for detecting the duration of LacZ expression after social defeat, we found a large number of LacZ cells in the CA3 region 1 day after social defeat. However, LacZ signals seemed to disappear quickly in the CA3 region so that almost no LacZ cells were found in the pyramidal layer of the CA3 region at 4 days after social defeat. Indeed, we saw high levels of cFos expression in the CA3 region during reactivation of CA1 and DG engram cells, suggesting the activation of CA3 cells during memory recall. It is unclear why long-term LacZ expression can be found in the CA1 and DG regions, but not in the CA3 region. Since LacZ expression is sustained by the tetracycline-insensitive tTA after the reintroduction of Dox food, the lack of CA3 LacZ signal may be due to poor expression of this mutated tTA in the CA3 region. The role of CA3 engrams in stress susceptibility cannot be ruled out, since CA3 engrams have been shown to be more sensitive to fear-related contextual information than neutral novel context (Denny et al., 2014). Using the Cre-dependent ArcCreER^T2^ mouse line may reveal the contribution of CA3 neurons to stress susceptibility.

**Figure 3:**
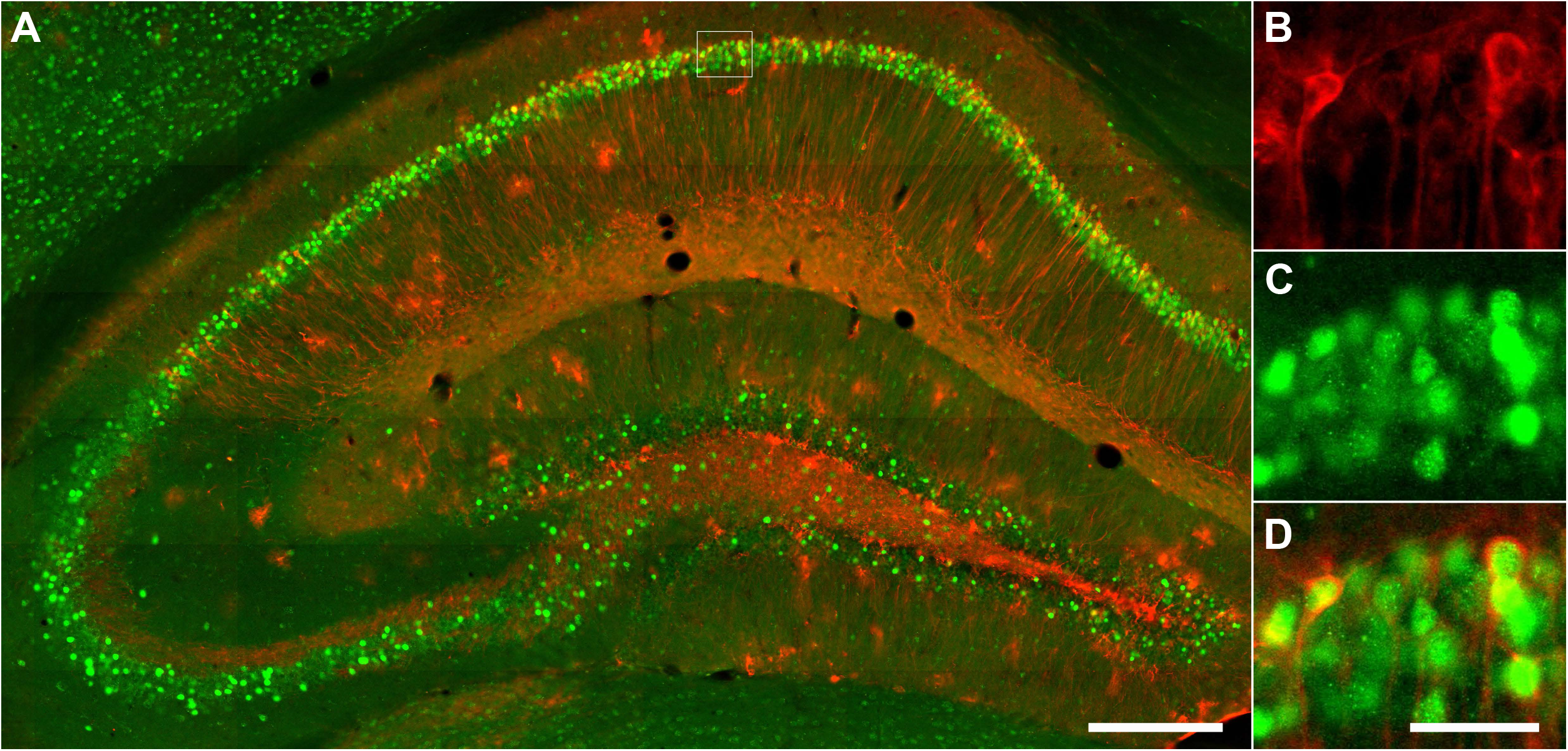
*Hippocampal LacZ and cFos staining of a TetTag mouse*. (**A**) Florescent micrographs of dorsal hippocampal CA1 neurons that were stained for LacZ (red) and cFos (green). Part of the CA1 (dotted line square) was enlarged to show LacZ (**B**), cFos (**C**) and the overlapping of LacZ and cFos in engram cells (**D**). Scale bars = 250 μm (**A**) and 40 μm (**B** to **D**).

The difference in negative memory engrams between susceptible and resilient mice have important implications for depression. Changes in these memory functions could be related to the bottom up changes from a hypersensitive medial temporal lobe, including hyperfunctioning of the amygdala and the hippocampus. Our findings that negative memory engrams are found in mice that are susceptible to social defeat stress suggested that these engrams could mechanistically contribute to the negative bias of memory formation in depression. Negative memory engrams correlated with the expression of social avoidance, suggesting their roles in mediating cognitive symptoms of depression. Inhibiting negative memory engrams in the hippocampus could be a novel therapeutic approach for treating cognitive symptoms in depression.

**Figure 4:**
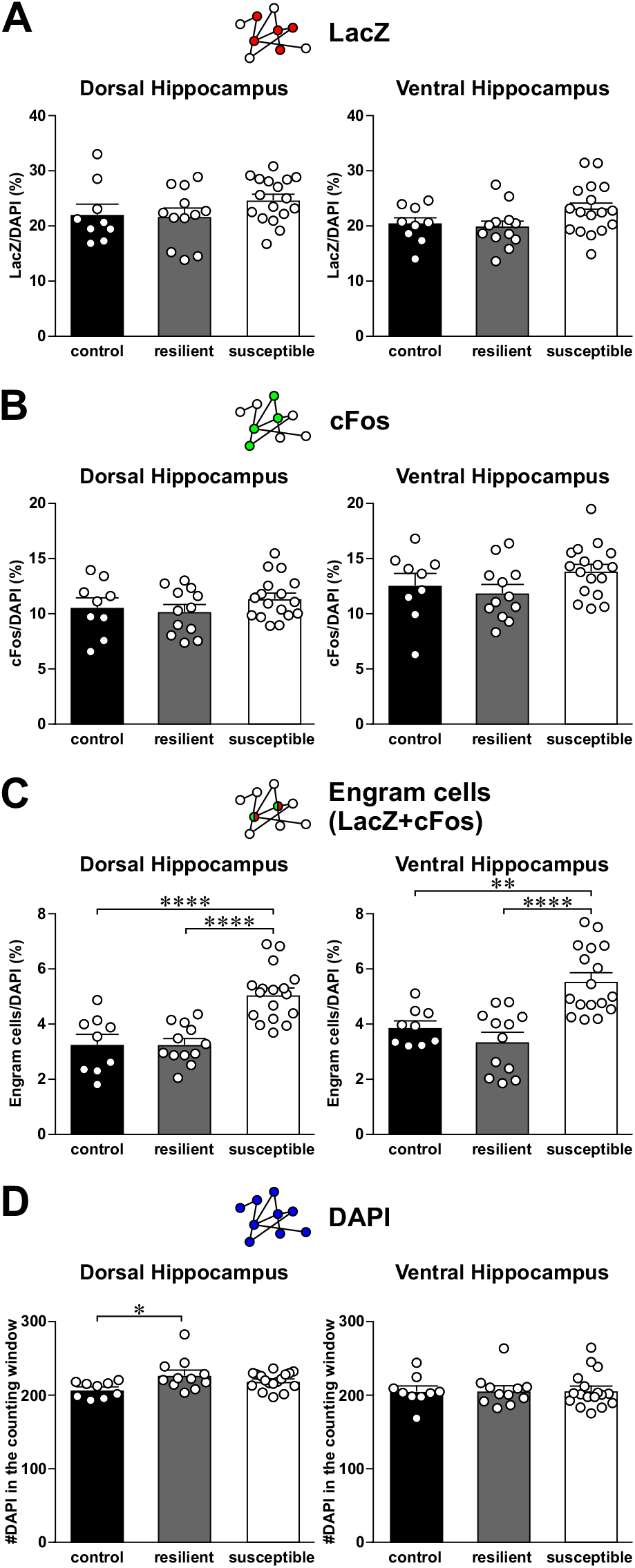
*Expression of LacZ, cFos and engram cells in the CA1 region of the dorsal and ventral hippocampus of control, resilient and susceptible mice*. (**A**) Histograms show the density of LacZ cells in the CA1 region of dorsal (left) and ventral hippocampus (*right*) of control, resilient and susceptible mice. (**B**) Histograms show the density of cFos cells in the CA1 region of dorsal (*left*) and ventral hippocampus (*right*) of control, resilient and susceptible mice. (**C**) Histograms show the density of engram cells (double labeled for both LacZ and cFos) in the CA1 region of dorsal (*left*) and ventral hippocampus (*right*) of control, resilient and susceptible mice. ** p < 0.01, **** p < 0.0001, Tukey’s test after ANOVA. (**D**) Histograms show the density of DAPI cells in the CA1 region of dorsal (left) and ventral hippocampus (*right*) of control, resilient and susceptible mice. * p < 0.05, Tukey’s test after ANOVA.

**Figure 5:**
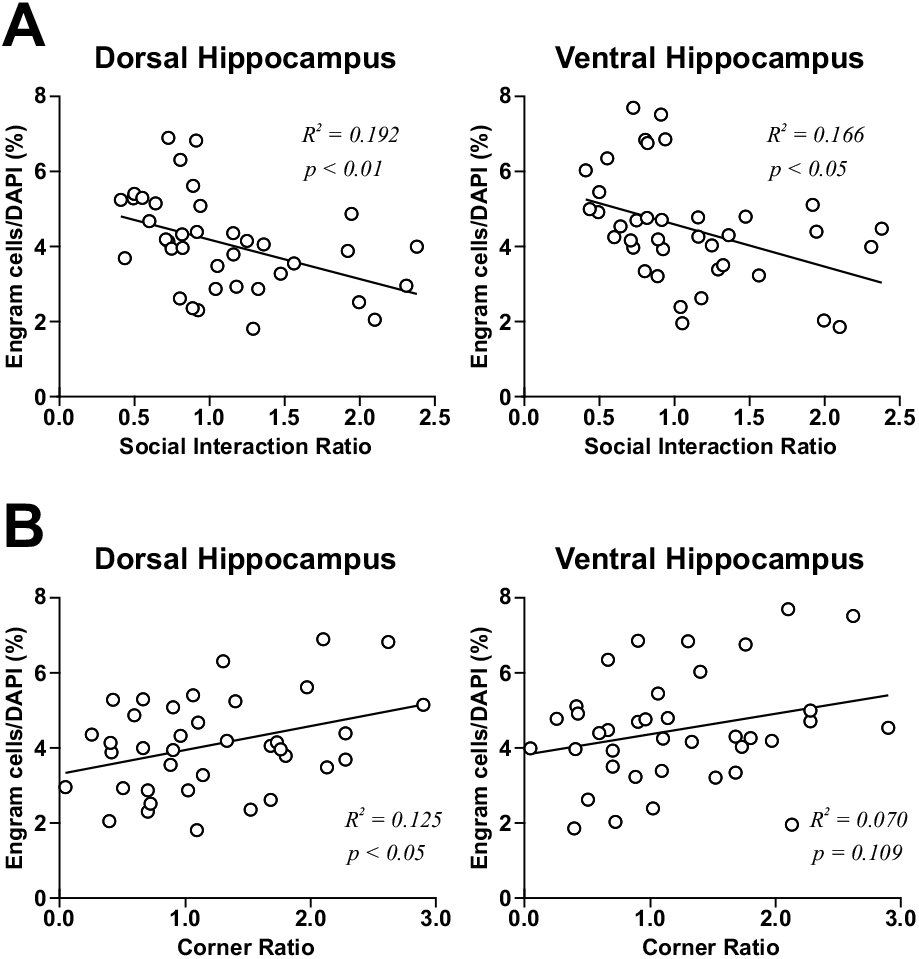
*Density of CA1 engram cells correlates with depression-related behaviors*. (**A**) Scatter plots of engram cells density in the CA1 region of dorsal (*left*) and ventral hippocampus (*right*) vs. social interaction ratio of control and stressed mice. (**B**) Scatter plots of engram cells density in the CA1 region of dorsal (*left*) and ventral hippocampus (*right*) vs. corner ratio of control and stressed mice.

**Figure 6:**
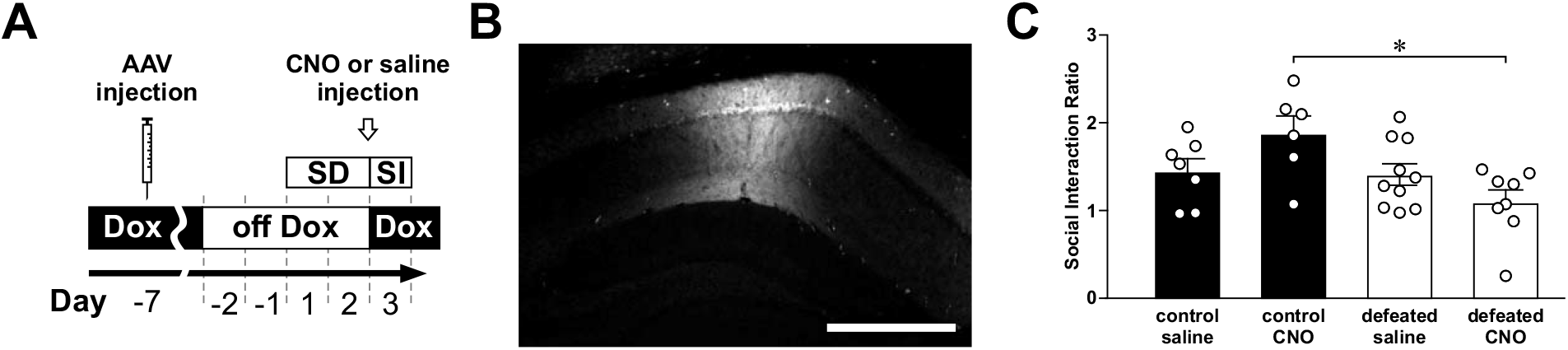
*Activation of social defeat-related CA1 engram cells reduces social interaction*. (**A**) A schematic diagram of the experimental design. cFos-tTA mice were bilaterally injected with AAV-PTRE-tight-hM3Dq-mCherry into the dorsal hippocampal CA1 region. One week later, they were off Dox for 2 days before being stressed by a subthreshold social defeat protocol, which consisted of two episodes of social defeat (SD). One day after defeat, mice were injected by either clozapine-N oxide (CNO, 3 mg/kg) or saline at 1 hour before the social interaction (SI) test. (**B**) A florescent micrograph shows the expression of AAV-PTRE-tight-hM3Dq-mCherry in the dorsal hippocampal CA1 region. Scale bar = 300 μm. (**C**) Histograms show the social interaction ratio of mice from different groups. * p < 0.05, Tukey’s test after ANOVA.

**Figure 7:**
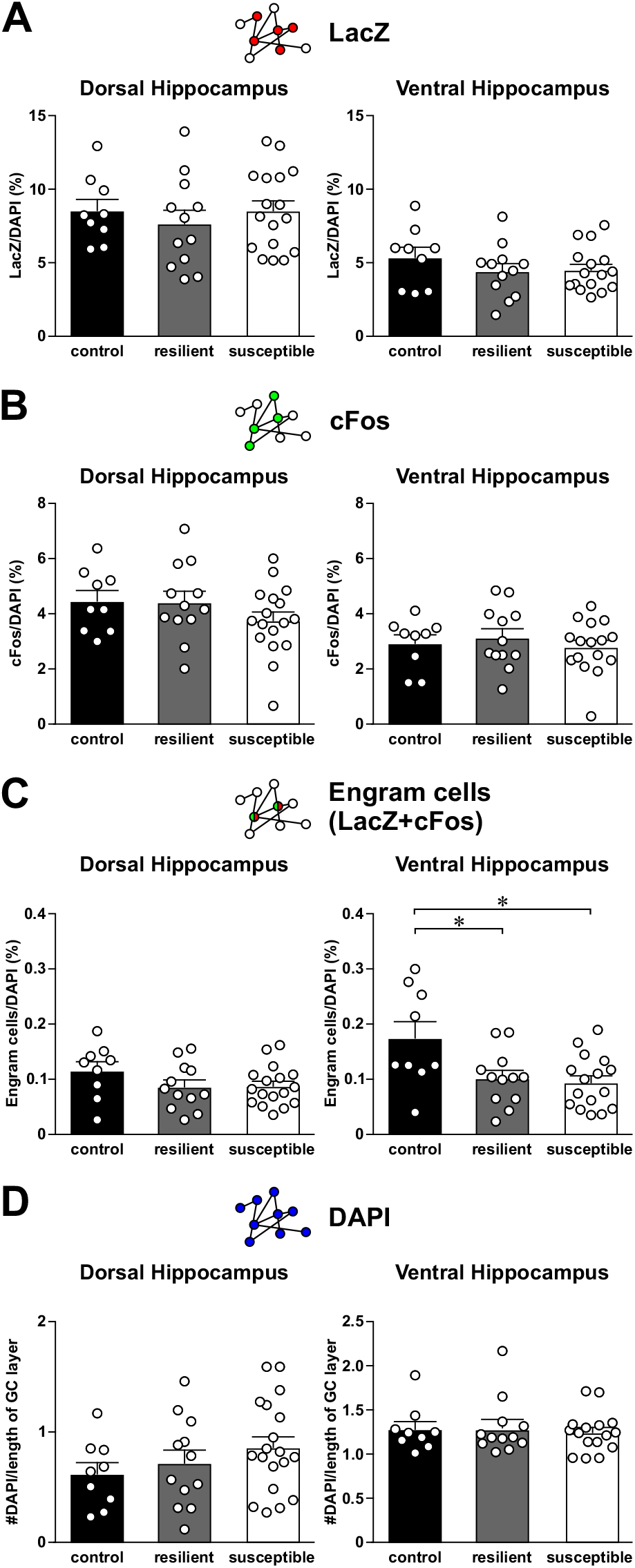
*Expression of LacZ, cFos and engram cells in the dental gyrus region of the dorsal and ventral hippocampus of control, resilient and susceptible mice*. (**A**) Histograms show the density of LacZ cells in the CA1 region of dorsal (left) and ventral hippocampus (*right*) of control, resilient and susceptible mice. (**B**) Histograms show the density of cFos cells in the CA1 region of dorsal (*left*) and ventral hippocampus (*right*) of control, resilient and susceptible mice. (**C**) Histograms show the density of engram cells (double labeled for both LacZ and cFos) in the CA1 region of dorsal (*left*) and ventral hippocampus (*right*) of control, resilient and susceptible mice. * p < 0.05, Tukey’s test after ANOVA. (**D**) Histograms show the density of DAPI (double labeled for both LacZ and cFos) stained cells in the CA1 region of dorsal (*left*) and ventral hippocampus (*right*) of control, resilient and susceptible mice.

**Figure 8:**
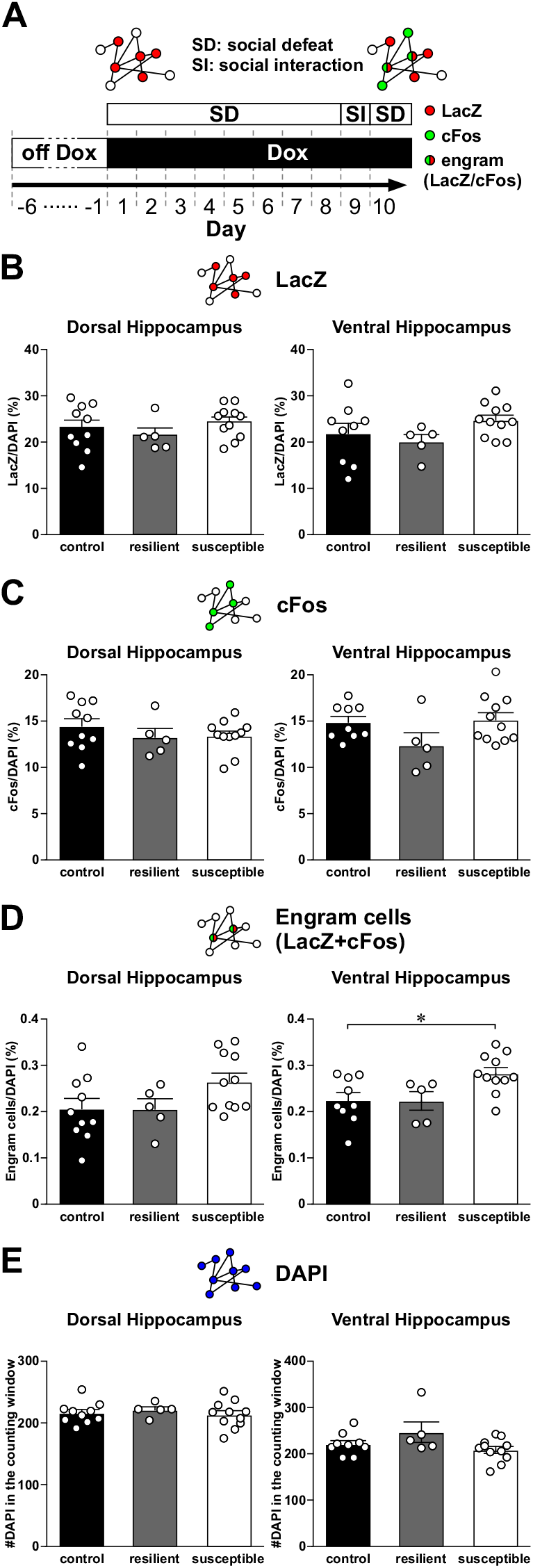
*Expression of LacZ, cFos and engram cells in the CA1 region of the dorsal and ventral hippocampus of control, resilient and susceptible mice, when labeling was stopped before social defeat*. (**A**) A schematic diagram of the experimental design. Tet-Tag mice were off Dox for 4 days. Labeling of neurons was stopped a day before the beginning of social defeat by feeding TetTag mice with doxycycline-containing food. Mice were then stressed by 8 episodes of social defeat (SD). The interaction between TetTag mice and a CD1 mouse, the strain of aggressive mice used for SD, was examined in a social (SI) interaction test. One day after the SI test, mice underwent one more episode of SD to trigger neuronal activation. Mice were sacrificed 90 minutes after the last episode of social defeat. Cartoons above the experimental plan depict the labeling of activated neurons during the first two days of chronic SD (red, LacZ), during the last episode of SD (green, cFos), and engram cells that expressed both signals (red/green). (**B**) Histograms show the density of LacZ cells in the CA1 region of dorsal (*left*) and ventral hippocampus (*right*) of control, resilient and susceptible mice. (**C**) Histograms show the density of cFos cells in the CA1 region of dorsal (left) and ventral hippocampus (*right*) of control, resilient and susceptible mice. (**D**) Histograms show the density of engram cells (double labeled for both LacZ and cFos) in the CA1 region of dorsal (*left*) and ventral hippocampus (*right*) of control, resilient and susceptible mice. * p < 0.05, Tukey’s test after ANOVA. (**E**) Histograms show the density of DAPI cells in the CA1 region of dorsal (*left*) and ventral hippocampus (*right*) of control, resilient and susceptible mice.

**Figure 9:**
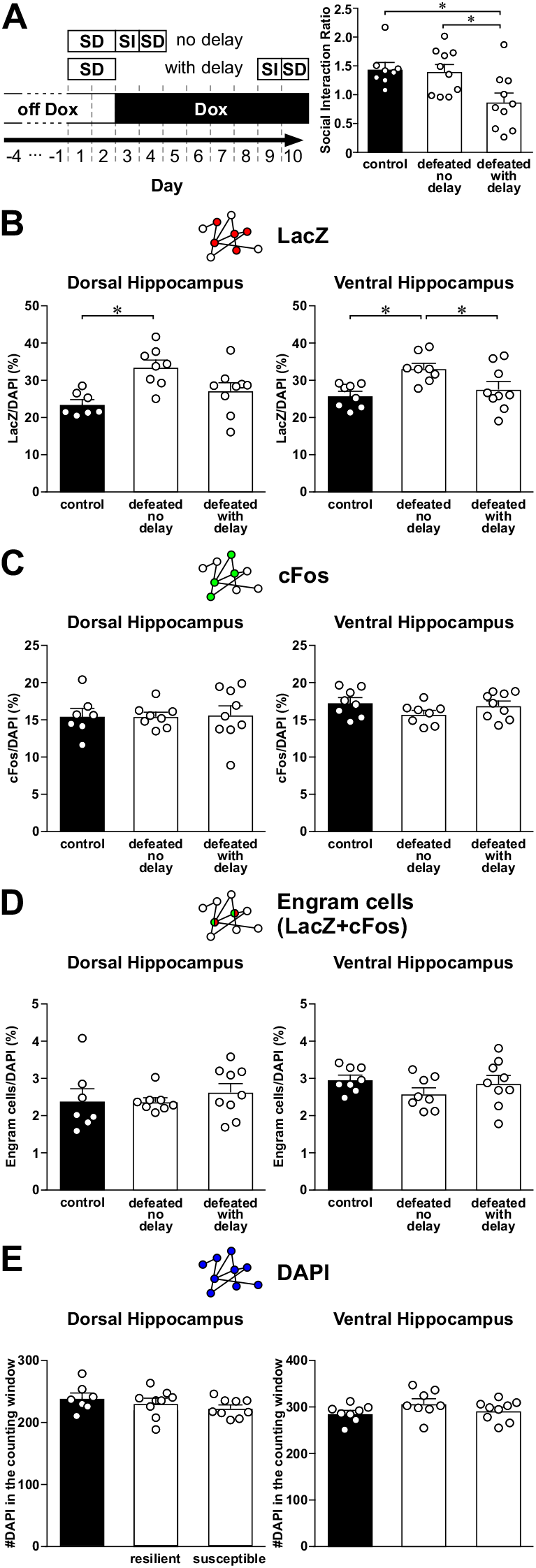
*Expression of LacZ, cFos and engram cells in the CA1 region of the dorsal and ventral hippocampus of mice that were stressed by a subthreshold social defeat protocol*. (**A**) *Left*: A schematic diagram of the experimental design. Tet-Tag mice were off Dox for 4 days. Mice were then stressed by 2 episodes of social defeat (SD). Labeling was blocked by putting mice on Dox-containing food after defeat. The interaction between TetTag mice and a CD1 mouse was examined in a social (SI) interaction test on 1 day (no delay) or 7 days (with delay) after SD. One day after the SI test, mice underwent one more episode of SD to trigger neuronal activation. Mice were sacrificed 90 minutes after the last episode of social defeat. *Right*: Histograms show the social interaction ratio of non-stressed control and defeated mice when SI tests were done either 1 day (no delay) or 7 days (with delay) after SD. * p < 0.05, Tukey’s test after ANOVA. (**B**) Histograms show the density of LacZ cells in the CA1 region of dorsal (*left*) and ventral hippocampus (*right*) of nonstressed control and defeated mice when SI tests were done either 1 day (no delay) or 7 days (with delay) after SD. * p < 0.05, Tukey’s test after ANOVA. (C) Histograms show the density of cFos cells in the CA1 region of dorsal (*left*) and ventral hippocampus (*right*) of non-stressed control and defeated mice when SI tests were done either 1 day (no delay) or 7 days (with delay) after SD. (**D**) Histograms show the density of engram cells (double labeled for both LacZ and cFos) in the CA1 region of dorsal (left) and ventral hippocampus (*right*) of non-stressed control and defeated mice when SI tests were done either 1 day (no delay) or 7 days (with delay) after SD. (**E**) Histograms show the density of DAPI cells in the CA1 region of dorsal (*left*) and ventral hippocampus (*right*) of non-stressed control and defeated mice when SI tests were done either 1 day (no delay) or 7 days (with delay) after SD.

## References

Abela JR, Hankin BL (2011) Rumination as a vulnerability factor to depression during the transition from early to middle adolescence: a multiwave longitudinal study. J Abnorm Psychol 120:259–271.

Alloy LB, Abramson LY, Whitehouse WG, Hogan ME, Tashman NA, Steinberg DL, Rose DT, Donovan P (1999) Depressogenic cognitive styles: predictive validity, information processing and personality characteristics, and developmental origins. Behav Res Ther 37:503–531.

Anacker C, Luna VM, Stevens GS, Millette A, Shores R, Jimenez JC, Chen B, Hen R (2018) Hippocampal neurogenesis confers stress resilience by inhibiting the ventral dentate gyrus. Nature 559:98–102.

Bagot RC, Parise EM, Pena CJ, Zhang HX, Maze I, Chaudhury D, Persaud B, Cachope R, Bolanos-Guzman CA, Cheer JF, Deisseroth K, Han MH, Nestler EJ (2015) Ventral hippocampal afferents to the nucleus accumbens regulate susceptibility to depression. Nat Commun 6:7062.

Botta P, Demmou L, Kasugai Y, Markovic M, Xu C, Fadok JP, Lu T, Poe MM, Xu L, Cook JM, Rudolph U, Sah P, Ferraguti F, Luthi A (2015) Regulating anxiety with extrasynaptic inhibition. Nat Neurosci 18:1493–1500.

Cai DJ et al. (2016) A shared neural ensemble links distinct contextual memories encoded close in time. Nature 534:115–118.

Conway MA, Pleydell-Pearce CW (2000) The construction of autobiographical memories in the self-memory system. Psychol Rev 107:261–288.

Deng W, Mayford M, Gage FH (2013) Selection of distinct populations of dentate granule cells in response to inputs as a mechanism for pattern separation in mice. Elife 2:e00312.

Denny CA, Kheirbek MA, Alba EL, Tanaka KF, Brachman RA, Laughman KB, Tomm NK, Turi GF, Losonczy A, Hen R (2014) Hippocampal memory traces are differentially modulated by experience, time, and adult neurogenesis. Neuron 83:189–201.

Denson TF, Pedersen WC, Ronquillo J, Nandy AS (2009) The angry brain: neural correlates of anger, angry rumination, and aggressive personality. J Cogn Neurosci 21:734–744.

Disner SG, Beevers CG, Haigh EA, Beck AT (2011) Neural mechanisms of the cognitive model of depression. Nat Rev Neurosci 12:467–477.

El Iskandrani KS, Oosterhof CA, El Mansari M, Blier P (2015) Impact of subanesthetic doses of ketamine on AMPA-mediated responses in rats: An in vivo electrophysiological study on monoaminergic and glutamatergic neurons. J Psychopharmacol 29:792–801.

Fanselow MS, Dong HW (2010) Are the dorsal and ventral hippocampus functionally distinct structures? Neuron 65:7–19.

Fu CH, Williams SC, Cleare AJ, Brammer MJ, Walsh ND, Kim J, Andrew CM, Pich EM, Williams PM, Reed LJ, Mitterschiffthaler MT, Suckling J, Bullmore ET (2004) Attenuation of the neural response to sad faces in major depression by antidepressant treatment: a prospective, event-related functional magnetic resonance imaging study. Arch Gen Psychiatry 61:877–889.

Gaddy MA, Ingram RE (2014) A meta-analytic review of mood-congruent implicit memory in depressed mood. Clin Psychol Rev 34:402–416.

Hamilton JP, Gotlib IH (2008) Neural substrates of increased memory sensitivity for negative stimuli in major depression. Biol Psychiatry 63:1155–1162.

Hertel PT, Gerstle M (2003) Depressive deficits in forgetting. Psychol Sci 14:573–578.

Joormann J, Quinn ME (2014) Cognitive processes and emotion regulation in depression. Depress Anxiety 31:308–315.

Josselyn SA, Kohler S, Frankland PW (2015) Finding the engram. Nat Rev Neurosci 16:521–534.

Kircanski K, Joormann J, Gotlib IH (2012) Cognitive Aspects of Depression. Wiley Interdiscip Rev Cogn Sci 3:301–313.

Koster EH, De Raedt R, Leyman L, De Lissnyder E (2010) Mood-congruent attention and memory bias in dysphoria: Exploring the coherence among information-processing biases. Behav Res Ther 48:219–225.

Krishnan V et al. (2007) Molecular adaptations underlying susceptibility and resistance to social defeat in brain reward regions. Cell 131:391–404.

Leutgeb JK, Leutgeb S, Moser MB, Moser EI (2007) Pattern separation in the dentate gyrus and CA3 of the hippocampus. Science 315:961–966.

Liu X, Ramirez S, Pang PT, Puryear CB, Govindarajan A, Deisseroth K, Tonegawa S (2012) Optogenetic stimulation of a hippocampal engram activates fear memory recall. Nature 484:381–385.

Lyubomirsky S, Caldwell ND, Nolen-Hoeksema S (1998) Effects of ruminative and distracting responses to depressed mood on retrieval of autobiographical memories. J Pers Soc Psychol 75:166–177.

Maeng S, Zarate CA, Jr., Du J, Schloesser RJ, McCammon J, Chen G, Manji HK (2008) Cellular mechanisms underlying the antidepressant effects of ketamine: role of alpha-amino-3-hydroxy-5-methylisoxazole-4-propionic acid receptors. Biol Psychiatry 63:349–352.

Mandell D, Siegle GJ, Shutt L, Feldmiller J, Thase ME (2014) Neural substrates of trait ruminations in depression. J Abnorm Psychol 123:35–48.

Mayberg HS, Brannan SK, Tekell JL, Silva JA, Mahurin RK, McGinnis S, Jerabek PA (2000) Regional metabolic effects of fluoxetine in major depression: Serial changes and relationship to clinical response. Biol Psychiatry 48:830–843.

McEwen BS (2004) Protection and damage from acute and chronic stress: allostasis and allostatic overload and relevance to the pathophysiology of psychiatric disorders. Ann N Y Acad Sci 1032:1–7.

McGonagle KA, Kessler RC (1990) Chronic stress, acute stress, and depressive symptoms. Am J Community Psychol 18:681–706.

McKinnon MC, Yucel K, Nazarov A, MacQueen GM (2009) A meta-analysis examining clinical predictors of hippocampal volume in patients with major depressive disorder. J Psychiatry Neurosci 34:41–54.

Nolen-Hoeksema S (2000) The role of rumination in depressive disorders and mixed anxiety/depressive symptoms. J Abnorm Psychol 109:504–511.

Radulovic J, Kammermeier J, Spiess J (1998) Relationship between fos production and classical fear conditioning: effects of novelty, latent inhibition, and unconditioned stimulus preexposure. J Neurosci 18:7452–7461.

Ramirez S, Liu X, MacDonald CJ, Moffa A, Zhou J, Redondo RL, Tonegawa S (2015) Activating positive memory engrams suppresses depression-like behaviour. Nature 522:335–339.

Reijmers LG, Perkins BL, Matsuo N, Mayford M (2007) Localization of a stable neural correlate of associative memory. Science 317:1230–1233.

Roy DS, Kitamura T, Okuyama T, Ogawa SK, Sun C, Obata Y, Yoshiki A, Tonegawa S (2017) Distinct Neural Circuits for the Formation and Retrieval of Episodic Memories. Cell 170:1000–1012 e1019.

Rude SS, Valdez CR, Odom S, Ebrahimi A (2003) Negative cognitive biases predict subsequent depression. Cognit Ther Res 27:415–429.

Santarelli L, Saxe M, Gross C, Surget A, Battaglia F, Dulawa S, Weisstaub N, Lee J, Duman R, Arancio O, Belzung C, Hen R (2003) Requirement of hippocampal neurogenesis for the behavioral effects of antidepressants. Science 301:805–809.

Squire LR (1992) Memory and the hippocampus: a synthesis from findings with rats, monkeys, and humans. Psychol Rev 99:195–231.

Stefanelli T, Bertollini C, Luscher C, Muller D, Mendez P (2016) Hippocampal Somatostatin Interneurons Control the Size of Neuronal Memory Ensembles. Neuron 89:1074–1085.

Thomas EJ, Elliott R, McKie S, Arnone D, Downey D, Juhasz G, Deakin JF, Anderson IM (2011) Interaction between a history of depression and rumination on neural response to emotional faces. Psychol Med 41:1845–1855.

Tonegawa S, Liu X, Ramirez S, Redondo R (2015) Memory Engram Cells Have Come of Age. Neuron 87:918–931.

Tse YC, Montoya I, Wong AS, Mathieu A, Lissemore J, Lagace DC, Wong TP (2014) A longitudinal study of stress-induced hippocampal volume changes in mice that are susceptible or resilient to chronic social defeat. Hippocampus 24:1120–1128.

Venzala E, Garcia-Garcia AL, Elizalde N, Delagrange P, Tordera RM (2012) Chronic social defeat stress model: behavioral features, antidepressant action, and interaction with biological risk factors. Psychopharmacology (Berl) 224:313–325.

Videbech P, Ravnkilde B (2004) Hippocampal volume and depression: a meta-analysis of MRI studies. Am J Psychiatry 161:1957–1966.

Wiltgen BJ, Silva AJ (2007) Memory for context becomes less specific with time. Learn Mem 14:313–317.

Xu W, Sudhof TC (2013) A neural circuit for memory specificity and generalization. Science 339:1290–1295.

Yokoyama M, Matsuo N (2016) Loss of Ensemble Segregation in Dentate Gyrus, but not in Somatosensory Cortex, during Contextual Fear Memory Generalization. Front Behav Neurosci 10:218.

Zhang Z, Ferretti V, Guntan I, Moro A, Steinberg EA, Ye Z, Zecharia AY, Yu X, Vyssotski AL, Brickley SG, Yustos R, Pillidge ZE, Harding EC, Wisden W, Franks NP (2015) Neuronal ensembles sufficient for recovery sleep and the sedative actions of alpha2 adrenergic agonists. Nat Neurosci 18:553–561.

